# Proteomic insights into mental health status: plasma markers in young adults

**DOI:** 10.1101/2023.06.07.544039

**Authors:** Alexey M. Afonin, Aino-Kaisa Piironen, Izaque de Sousa Maciel, Mariia Ivanova, Arto Alatalo, Alyce M Whipp, Lea Pulkkinen, Richard J Rose, Irene van Kamp, Jaakko Kaprio, Katja M. Kanninen

## Abstract

Global emphasis on enhancing prevention and treatment strategies necessitates increased understanding of biological mechanisms of psychopathology. Plasma proteomics is a powerful tool that has been applied in the context of specific mental disorders for biomarker identification. The p-factor, also known as the “general psychopathology factor”, is a concept in psychopathology suggesting that there is a common underlying factor that contributes to the development of various forms of mental disorders. It has been proposed that the p-factor can be used to understand the overall mental health status of an individual. Here we aimed to discover plasma proteins associated with the p-factor in 775 young adults in the FinnTwin12 cohort. Using liquid chromatography–tandem mass spectrometry, 14 proteins with a significant connection with the p-factor were identified, 9 of which were linked to epidermal growth factor receptor (EGFR) signalling. This exploratory study provides new insight into biological alterations associated with mental health status in young adults.

## Introduction

Mental health issues are increasingly becoming a major concern globally (Organization 2022; World Health Organization 2021). In fact, the World Health Organization estimates that about one in every eight people across the globe suffer from a mental health disorder, making these disorders the primary cause of a reduced quality of life (Organization 2022). The recent COVID-19 pandemic has notably exacerbated mental health issues, particularly among young adults aged 18–29. During the pandemic, the number of young adults experiencing depression symptoms more than doubled in numerous European countries (*Health at a Glance: Europe 2022* 2022).

Despite the significant impact of mental health diseases on daily life and their considerable economic cost, these conditions often go undiagnosed and untreated (*Health at a Glance: Europe 2022* 2022; Organization 2022). This highlights the pressing need for early detection of individuals at high risk for psychopathology, targeted preventative measures, and improvements in diagnostic procedures and treatments.

Although different mental disorders may have unique symptoms, they have been shown to share commonalities in terms of underlying biological, psychological, and social factors (Smith et al. 2020). The p-factor, also known as the “general psychopathology factor,” is a concept in psychopathology suggesting that there is a common underlying factor that contributes to the development of various forms of mental disorders (Caspi et al. 2014; Lahey et al. 2012). It has been proposed that this single latent factor can encapsulate individuals’ proclivity to develop all forms of psychopathology included within the broad internalising, externalising and thought disorder dimensions (Ronald 2019). The p-factor is analogous to the general factor in intelligence (called the g-factor) which summarizes the observation that individuals who do well on one type of cognitive test tend to do well on all other types of cognitive tests (Caspi et al. 2014; Jensen 1999). Other factors, such as general factor of personality (GFP) and a general factor of personality disorder (g-PD) have been previously shown to have high correlation with the p-factor (Oltmanns et al. 2018). At the individual level, the p-factor reflects meaningful differences between persons on a single dimension that represents the tendency to experience psychiatric problems as persistent and comorbid; that is, high p-factor individuals experience difficulties in regulation/control when dealing with others, the environment, and the self(Caspi et al. 2014; Laceulle, Vollebergh, and Ormel 2015; Smith et al. 2020).

Previous studies have shown the p-factor to be connected to brain functioning in adolescents, with higher p-factor scores associated with diminished activation of multiple brain zones during executive tasks (Shanmugan et al. 2016). Importantly, some studies have reported that the p-factor may be a stronger predictor of mental health outcomes than specific diagnoses of mental disorders (Hartwig et al. 2017). A recent study showed that the p-factor was associated with poorer performance on the simple reaction time task and the inspection time task, with speed of processing being a common correlate of psychopathology factors (Haywood et al. 2022). Likewise, Pulkkinen (Pulkkinen 2017) has shown that low emotion and behavior regulation observed as externalizing and internalizing problems in children are negatively associated with the executive functions of the forebrain for inhibition and updating (containing working memory and shifting). This suggests that the p-factor could be used to better understand the overall mental health status of an individual, rather than just focusing on individual diagnoses.

Biomarker discovery has gained traction in recent years as researchers seek to uncover the biological underpinnings of mental health conditions (Walters et al. 2018; Munn-Chernoff et al. 2021). The development of “omics” technologies and state-of-the-art analytical methods have increased interest in the capabilities of plasma proteomics in biomarker discovery. LC-MS/MS-based proteomics provides a global snapshot of protein expression patterns that reflect physiological and pathological states (Orlando and Aebersold 2019), making comprehensive analysis of the plasma proteome possible (Zhou et al. 2020). This has enabled the simultaneous detection and quantification of thousands of proteins, expediting biomarker discovery efforts and reducing the time and resources required for this process. This holistic view of proteomics allows for the unbiased discovery of novel biomarkers, with less need for prior knowledge of target proteins. This feature is particularly important in cases where the biology of the process is not yet fully understood or when new, unforeseen biomarkers are needed for improved diagnostic or prognostic applications (Ignjatovic et al. 2019).

Proteomics approaches have been utilized for the identification of protein signatures associated with specific psychological disorders (Comes et al. 2018; Domenici et al. 2010; Ziani et al. 2022). For example, several growth factors (BDNF, VEGF, NGF) and cytokines (IL-1β, IL-6, IFN-α) have been linked to depression (Malik et al. 2021). Moreover, a recent multi-omics study reported reduced apolipoprotein levels and an increase in complement effector proteins in the plasma of schizophrenia (SCZ) patients (Campeau et al. 2022). However, proteomics analyses have not previously been combined to studies of the p-factor for identification of markers associated with overall mental health status.

The FinnTwin12 (FT12) cohort, a longitudinal study of Finnish twins born between 1983 and 1987, has a multitude of data and biological samples (Kaprio 2006; Rose et al. 2019). As a valuable resource for exploring biological processes involved in mental health problems, we explored the connection between the p-factor and plasma proteomics among young adults from this cohort.

## Methods

### Cohort description

The FT12 cohort is a longitudinal population-based cohort of Finnish twins born 1983–1987 collected to investigate behavioural development and health habits (Kaprio 2006; Rose et al. 2019). Initially, twins and their families were identified using the Finnish Central Population Registry, and questionnaire collection occurred for all participants in the cohort at ages 11/12, 14, 17, and 22. The baseline response rate was 87% (N=5600 twins) and has remained high (response rate range: 85–90%). At age 14, a subset of the twins (from 1035 families) was more intensively studied, including psychiatric interviews and additional questionnaires (ages 14 and 22), as well as blood plasma samples (age 22). The “age 22” assessment wave of these more intensively studied twins involved 1347 individuals (mean age = 22.4 years, SD = 0.70; response rate 73.0%), 779 of whom attended in-person assessments and provided venous blood plasma samples. The blood samples were collected after overnight fasting, which involved abstaining from alcohol and tobacco since the night before sampling. Plasma was immediately extracted and stored at −80°C (Whipp et al. 2022).

### p-factor calculation

In FT12, behavioural and emotional characteristics were measured at all data collection waves. The modified Multidimensional Peer Nomination Inventory (MPNI) measure aimed at observing individual differences in emotion and behaviour regulation was used. It is an extension of the measure (Pulkkinen, Kaprio, and Rose 1999) used in the Jyväskylä Longitudinal Study of Personality and Social Development in which the development of the same individuals has been followed from age 8 to 50, with findings that low self-regulation is associated with social and psychological dysfunction (Pulkkinen 2017). The MPNI was collected in FT12 at ages 12, 14, and 17, from different raters (7 in total): parents (age 12), teachers (age 12 and 14), twin children themselves (age 14 and 17) and the child’s co-twin (age 14 and 17). The measure includes subscales of the externalizing problem dimension: aggression (6 items [for MPNI ages 12, 14, 17]), hyperactivity-impulsivity (7 items [MPNI ages 12, 14], 6 items [MPNI ages 17]), and inattention (4 items [MPNI ages 12, 14, 17]), as well as for the internalizing problem dimension; depression (5 items [MPNI ages 12, 14], 2 items [MPNI ages 17]), social anxiety (2 items [MPNI ages 12, 14], 3 items [MPNI ages 17]), and 1 item for victimization (MPNI ages 12, 14, 17). Each MPNI item (e.g., “Is restless, unable to sit still”) has four response choices (from “not observed in the child” to “clearly observable in the child”, scored 0–3 respectively). The MPNI p-factor score was created by combining all the items of the “externalising” and “internalising” dimensions together into a sum score, with at most 2 missing items allowed. Missing items were imputed based on the mean of the remaining items, with less than 3% of twins having missing items. A composite “combined” p-factor score was created using the p-factor scores of all seven of the abovementioned available MPNI ratings (Cronbach’s alpha=0.76), because we know that ratings from different raters are not highly correlated, however, they can impart unique information (Achenbach et al., 1987; Whipp et al., 2019, 2021). Each of the seven scores were standardized as z-scores, then we took the mean of available scores. Eleven twins had no overall p-factor score, leaving 775 twins. Of them, 505 (65%) had been scored by all raters at all times, while 194 (25%) had only one rater value missing, the remaining 10% having scores from 2–4 raters.

### High-abundance protein depletion

Albumin accounts for 50% and the top 22 proteins account for 99% of plasma proteins by weight in human plasma samples (Ignjatovic et al., 2019). Therefore, the depletion of high-abundant proteins is essential to the identification and analysis of low-abundant proteins. A commercial kit (High Select™ Top14 Abundant Protein Depletion Mini Spin Columns, cat. Number: A36370, ThermoScientific) was used to deplete the 14 most abundant proteins from plasma before the proteomic analyses. The depleted proteins were human serum albumin (HSA), albumin, IgG, IgA, IgM, IgD, IgE, kappa and lambda light chains, alpha-1-acidglycoprotein, alpha-1-antitrypsin, alpha-2-macroglobulin, apolipoprotein A1, fibrinogen, haptoglobin, and transferrin, according to manufacturer’s manual. Briefly, 10 µL of total plasma was added to the mini spin columns and incubated for 10 min while rotating, followed by centrifugation of the columns (1,000 *x g*) for 2 min. The filtrate was collected in 2 ml plastic tubes and stored at −20°C until preparation for mass spectrometry proteomic analyses, which were performed at the Turku Proteomics Facility in Finland supported by Biocenter Finland.

### Protein precipitation and digestion

The proteins of 786 depleted plasma samples were acetone precipitated and subjected to in-solution digestion according to standard protocol at the Turku Proteomics Facility, Turku, Finland (https://bioscience.fi/). After digestion, peptides were desalted with a Sep-Pak C18 96-well plate (Waters), evaporated to dryness, and stored at −20 ⁰C.

### Mass spectrometry analysis

Digested peptide samples were dissolved in 0.1% formic acid, and the peptide concentration was determined with a NanoDrop device. For data independent acquisition (DIA) analysis, 500 ng of peptides were injected and analysed in a random order. Wash runs were submitted between each sample to reduce potential carry-over of peptides. The Liquid Chromatography-Electrospray Ionization-Mass Spectrometry (LC-ESI-MS/MS) analysis was performed on a nanoflow HPLC system (Easy-nLC1000, Thermo Fisher Scientific) coupled to a Q Exactive HF mass spectrometer (Thermo Fisher Scientific, Bremen, Germany) equipped with a nano-electrospray ionization source. Peptides were first loaded on a trapping column and subsequently separated inline on a 15 cm C18 column (75 μm x 15 cm, ReproSilPur 3 μm 120 Å C18-AQ, Dr. Maisch HPLC GmbH, Ammerbuch-Entringen, Germany). The mobile phase consisted of water with 0.1% formic acid (solvent A) or acetonitrile/water (80:20 (v/v)) with 0.1% formic acid (solvent B). A 50 min from 5% to 35% solvent B gradient was used to elute peptides. Samples were analysed by a DIA LC-MS/MS method. MS data was acquired automatically by using Thermo Xcalibur 4.1 software (Thermo Fisher Scientific). In the DIA method, a duty cycle contained one full scan (400-1000 m/z) and 25 DIA MS/MS scans covering the mass range 400–1000 with variable width isolation windows.

### Protein identification and quantification analysis

Data analysis consisted of protein identification and label-free quantifications of protein abundances. Data was analysed by Spectronaut software (Biognosys; version 17.0.2). The direct DIA approach was used to identify proteins. The main data analysis parameters in Spectronaut were: (i) Enzyme: Trypsin/P; (ii) up to 2 missed cleavages; (iii) Fixed modifications: Carbamidomethyl (cysteine); (iv) Variable modifications: Acetyl (protein N-terminus) and oxidation (methionine); (v) Precursor FDR Cutoff: 0.01; (vi) Protein FDR Cutoff: 0.01; (vii) Quantification MS level: MS2; (viii) Quantification type: Area under the curve within integration boundaries for each targeted ion; (ix) Protein database: Homo sapiens Swiss-Prot reference proteome (Uniprot release 2022_01_07_HUMAN) (Bateman et al. 2023), Universal Protein Contaminant database (Frankenfield et al. 2022).

### Raw data drift and batch correction

The raw data investigation pre-processing and statistical analyses were performed in the R (version 4.2.1.) environment (R Core Team, 2022). The signal drift and the observed batch effect were corrected using the proBatch (v. 1.13.0) (Čuklina et al. 2021) package.

Protein abundances were analysed by LC-MS/MS in three separate experimental runs or batches. Since the number of samples in each batch was relatively large, the data was normalised before further analysis, and batch effects were removed. For the ease of comparing the LC-MS/MS runs, 10 of the samples were analysed in 2 out of the 3 runs. For the raw data analysis, we extracted the data from the .sne file using the iq export scheme (Pham et al., 2020). The data used for normalisation was the raw peak area of the peptide groups. These values were used in the further analyses.

The signal drift and the observed batch effect were corrected using the proBatch (v. 1.13.0) package (Čuklina et al. 2021). The median abundance plots showed the samples forming four distinct groups, identical to the batches of instrument runs (Supplementary figure 1A). The figure also shows pronounced signal drift in the third and fourth batches. These effects were corrected for using the proBatch pipeline. After the batch effect correction, no significant drift or batch effect could be seen (Supplementary figure 1B).

### Bioinformatic analysis

After drift and batch correction the fastMaxLFQ method from the iq package (v. 1.9.10) (Pham, Henneman, and Jimenez 2020) was used to transform the peptide abundancies into protein abundance values which were used for all the subsequent analyses. Only the identified proteins with quantified abundance levels in at least 80% of the samples were used in further analyses. Missing values remaining in the dataset were imputed using the Sample Minimum method (Liu and Dongre 2021). The connection between the p-factor and the protein abundances was analysed using the limma (Phipson et al. 2016) package (v. 3.54.2). Sex and age were included into linear models as covariates to ensure reported associations were not due to sex or age effects. Limma modelling was used to investigate the association of protein abundance with the p-factor using linear and non-linear modelling. The possible non-linear relationship between the p-factor and the protein abundance was investigated with using splines in limma (Ritchie et al. 2015). A basis matrix for representing the family of piecewise-cubic splines with 5 nodes were generated using the *ns* function from the p-factor variable (Splines package v 3.6.2), and was used in limma modelling, also including sex and age as covariates.

Moderate F-test on the p-factor was carried out to assess the significance of non-linear associations of the protein abundance with the p-factor using the function lmFit and eBayes from the R limma package. P values for linear and nonlinear modelling were corrected for multiple testing and the false discovery rate (FDR) was computed by using the Benjamini & Hochberg method (Benjamini and Hochberg 1995), which were reported as q-values. The significance level considered in all analyses was 0.05, unless stated otherwise. The linear effect size is reported as the log2-fold-change in expression that results from a unit (one standard deviation) change in p-factor.

The protein–protein interaction information for the significantly differentially abundant proteins was analysed using STRING database analysis (Szklarczyk et al. 2021). The enrichment analysis with Gene Ontology (Process, Function and Component), KEGG and Reactome pathways, PubMed publications, UniProt Keywords, and PFAM/INTERPRO/SMART domain databases was performed using the STRINGdb package (Szklarczyk et al. 2019). Result visualizations were performed using R and ggplot2 (v3.4.0) (Wickham 2009).

## Results

### Cohort characteristics

The p-factor was calculated based on assessments by multiple raters at three different ages as described in the Materials and Methods. A combined p-factor value was available for 775 individuals (318 males and 457 females). The z-score based p-factor distribution is presented in Figure 1.

**Figure 1.**
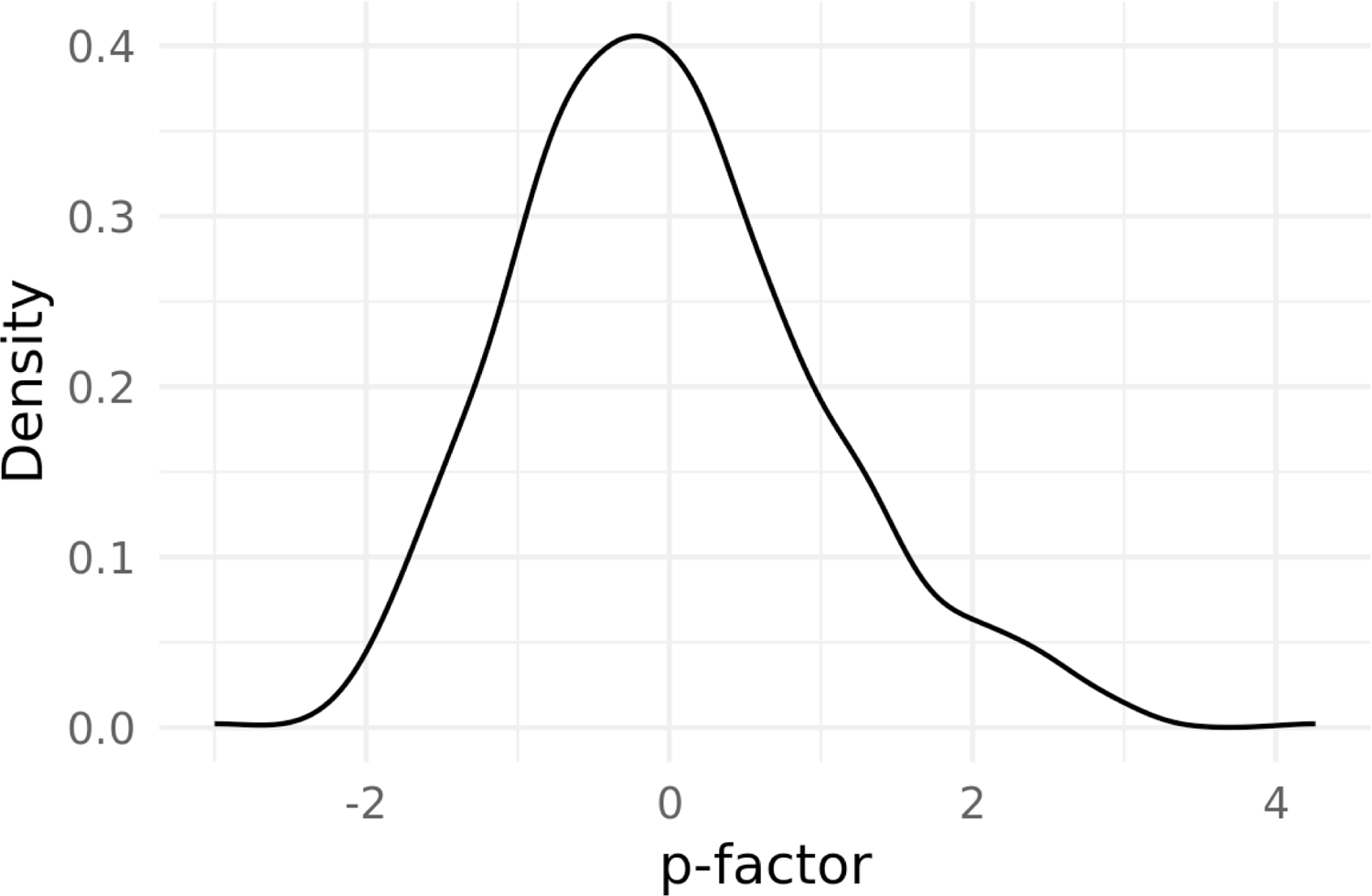
Density plot of the z-scored p-factor values of the participants in this study.

### Protein identification

MS-based proteomics successfully identified 1615 proteins (DIA spectrometry intensity values) in the FT12 cohort (N = 786). The mean number of identified proteins was 907 per sample (SD = 49). Only those proteins present in at least 80% of the samples were used for further analysis, leaving 636 proteins.

### Association of proteins with p-factor

The linear modelling showed 5 proteins inversely associated with the p-factor (Table 1). As the relationship between the altered proteins and p-factor is not known, the analysis was also performed using splines, which also made it possible to investigate non-linear relationships between the protein abundance and the p-factor. These twi analyses showed 14 proteins associated with the p-factor (Table 1). The relationships between the p-factor and the protein abundance for the significantly associated proteins are presented in Supplementary Figure 2.

**Table 1.** The plasma proteins significantly associated with the p-factor.

| *Protein ID* | *Gene Name* | *Protein name* | *Linear effect size* | *Linear q-value* | *Nonlinear q-value* |
| --- | --- | --- | --- | --- | --- |
| **Proteins with linear relationship with p-factor** |  |  |  |  |  |
| Q15828* | CST6 | Cystatin E/M | -0.086 | 0.024 | 0.047 |
| P98160 | HSPG2 | Heparan sulfate proteoglycan 2 | -0.061 | 0.006 | 0.066 |
| P23142 | FBLN1 | Fibulin 1 | -0.065 | 0.006 | 0.075 |
| P07911* | UMOD | Uromodulin | -0.145 | 0.006 | 0.013 |
| Q9Y6R7* | FCGBP | Fc gamma binding protein | -0.082 | 0.005 | 0.007 |
| **Proteins with nonlinear relationship with p-factor** |  |  |  |  |  |
| P04179 | SOD2 | Superoxide dismutase 2 | - | 0.701 | 0.001 |
| Q9BY67 | CADM1 | Cell adhesion molecule 1 | - | 0.329 | 0.002 |
| Q14126 | DSG2 | Desmoglein 2 | - | 0.072 | 0.010 |
| Q8NBJ4 | GOLM1 | Golgi membrane protein 1 | - | 0.866 | 0.010 |
| P07858 | CTSB | Cathepsin B | - | 0.496 | 0.010 |
| Q86UN3 | RTN4RL2 | Reticulon 4 receptor like 2 | - | 0.675 | 0.012 |
| O75636 | FCN3 | Ficolin 3 | - | 0.183 | 0.026 |
| P07942 | LAMB1 | Laminin subunit beta 1 | - | 0.183 | 0.035 |
| P26447 | S100A4 | S100 calcium binding protein A4 | - | 0.051 | 0.047 |

\* indicates proteins with both linear and non-linear relationship

### Functional enrichment and annotation

The STRING protein–protein interaction networks functional enrichment analysis showed two connected clusters of proteins with three proteins each: cystatin-M (CST6), uromodulin (UMOD), cathepsin B (CTSB), and laminin subunit beta-1 (LAMB1), basement membrane-specific heparan sulfate proteoglycan core protein (HSPG2), and fibulin-1 (FBLN1), shown in colour in Figure 2. Investigation of the first layer of the string network showed that nine of the significant proteins were linked specifically through the epidermal growth factor receptor (EGFR) and transthyretin (TTR) (Figure 2). Both proteins were among the 636 proteins we investigated, though the q-values were above the significance threshold (q-values for both EGFR and TTR were 0.074).

**Figure 2.**
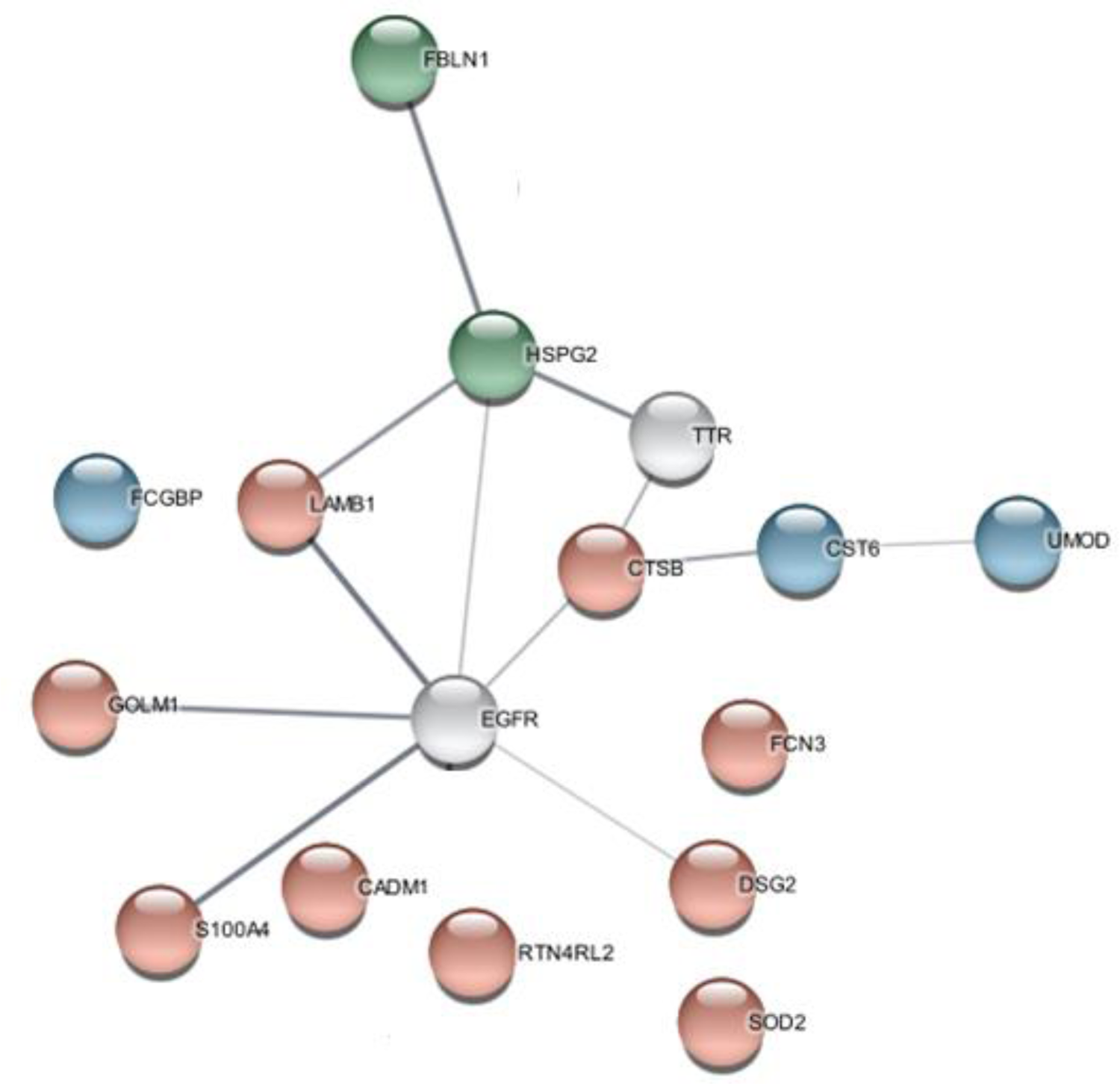
The result of STRING analysis of the proteins significantly associated with the p-factor. Line thickness indicates the strength of data support. Green circles denote proteins with a linear relationship to the p-factor, red circles with nonlinear relationships and blue circles representing proteins with reported linear and nonlinear relationships.

Enrichment analysis of function categories showed only the extracellular matrix structural constituent to be significantly enriched. Compartments, component, function, and tissue analyses showed significantly enriched terms, mostly connected to extracellular space and matrix, and cell–cell adhesion (Supplementary Table 1).

A connection to a disease of the CNS or other neurodegenerative disease according to the Disease Ontology database was found for 6 of the 14 significant proteins (Schriml et al. 2022), shown in Table 2.

**Table 2.** The disease ontology classification (STRINGdb)

| *Gene name* | *Disease ontology* |
| --- | --- |
| LAMB1, FCN3, CTSB, S100A4, HSPG2 | Nervous system disease |
| FCN3, CTSB, HSPG2, SOD2 | Neurodegenerative disease |
| FCN3, CTSB, S100A4, HSPG2 | Central nervous system disease |

## Discussion

The field of plasma proteomics is rapidly gaining traction in the realm of biomedical research, particularly in studies relating to mental health. There is a growing body of evidence to suggest that alterations in plasma protein profiles are associated with major psychiatric conditions, including major depressive disorder (MDD), schizophrenia, psychotic disorders (PSD), and bipolar disorders (BD) (Fernandes et al. 2022; García-Gutiérrez et al. 2020; Rodríguez Cerdeira et al. 2017). In this study, we present the first report of alterations to plasma proteins associated with the p-factor in young adults.

This study reports 14 plasma proteins associated with p-factor scores in young adults. All but the FCGBP protein were present in the Human Plasma Proteome Database (“Human Protein Atlas” 2022; Uhlén et al. 2019), FCGBP was also previously reported in serum samples (Ji et al. 2022). Of these proteins, nine belonged to a protein network connected to EGFR, six being directly connected to EGFR. The EGF-related signalling pathways have been previously linked to neurodevelopment (Birchmeier 2009), synaptic plasticity (Ledonne and Mercuri 2019; Mei and Nave 2014), chronic pain (Borges et al. 2021), fear (Chen et al. 2022), as well as to mental health diseases (Fernandes et al. 2022; Fiori et al. 2021; Mei and Nave 2014; Mei and Xiong 2008; Nawwar, Zaki, and Sayed 2022; Shi and Bergson 2020). For example, altered EGFR signalling has been reported in MDD and BD patients in blood proteomics studies (Fernandes et al. 2022).

In addition to EGFR signalling, we observed the p-factor to be negatively associated with heparan sulfate proteoglycan 2 (HSPG2). Heparan sulfate proteoglycans (HSPG) are membrane proteins and a major component of extracellular matrices involved in many cellular processes, as they function as co-receptors for growth factors (Bishop, Schuksz, and Esko 2007). HSPG2, combined with CEP350 and SMAD5, was recently presented as a potential diagnostic biomarker for MDD (Long et al. 2021). Furthermore, the *HSPG2* gene was previously connected to antipsychotic-induced adverse effects such as tardive dyskinesia (MacNeil and Müller 2016; Zai, Maes, et al. 2018), also specifically in SCZ patients (Zai, Lee, et al. 2018), and the maintenance and repair of the blood-brain barrier in mice (Nakamura et al. 2019). Moreover, downregulation of *HSPG2* and a depressive-like phenotype were revealed in mouse models of chronic mild stress and impaired glutamate function (Tordera et al. 2011). Taken together with the published evidence, our reported data suggests that HSPG2 has an important role in neuropsychology and mental health.

We also report a negative association with fibulin-1 (FBLN1) and the p-factor. The FBLN1 gene is connected to central nervous system development (Bohlega, Al-Ajlan, and Al-Saif 2014; Cooley et al. 2008) and modulation of neurotrophic activities of amyloid precursor protein in cultured rodent neural stem cells (Ohsawa, Takamura, and Kohsaka 2001). So far, little is known about the possible connection of FBLN1 to mental health. However, Shin *et al*. reported decreased FBLN1 plasma protein levels in MDD patients compared to BD patients and healthy controls, supporting our findings (Shin et al. 2021). The model proposed in their study also contained Fc gamma binding protein (FCGBP), which was reported to be significantly higher in BD patients compared to MDD patients but not in healthy controls. In our study, FCGBP was negatively associated with the p-factor. Additionally, increased plasma protein abundance of desmoglein 3 (DSG3) was reported in MDD patients and reduced abundance in BD patients compared to healthy controls (Shin et al. 2021). DSG3 is a protein belonging to the same desmosomal cadherin family as DSG2 reported in this study, which had a non-linear association with the p-factor with increased abundancies in the middle part of the p-factor scale. DSG2 was previously shown to have a similar function and was also shown to compensate for DSG3 in DSG3^-^ mouse models (Hartlieb et al. 2014). The Shin et al. paper investigated BD, which is classified as a Thought Disorder factor, and MDD, which is classified as an Internalising factor (Caspi et al. 2014). Since both the disorders can be included in the p-factor, which means the effects observed in their paper cannot be observed based on the p-factor alone.

Laminin subunit beta-1 (LAMB1) was shown to be connected to the p-factor in this study. A polymorphism in *LAMB1* gene has been earlier associated with autism severity (Kim et al. 2015), neural development of embryonic stem cells (Sun et al. 2008) and pain sensitivity in mice (Li et al. 2021). *LAMB1* is expressed during the early development of nervous system (Kim et al. 2015) and in the hippocampus in the mature brain (Yang et al. 2011). In rats, LAMB1 showed negative regulation of spatial learning through the inhibition of the ERK/MAPK-SGK1 signalling pathway in the hippocampus (Yang et al. 2011). Furthermore, loss of LAMB1 in the anterior cingulate cortex was found to increase pain sensitivity and be associated to anxiety- and depressive-like behaviour in mice (Li et al. 2021).

Cathepsin B (CTSB) was identified here with a nonlinear relationship to the p-factor. We also found two other significant proteins, CST6 and UMOD, which formed a cluster with CTSB according to the STRING database analysis. Moon *et al*. suggested CTSB as a mediator of exercise-induced effects on brain health by enhancing expression of neurotrophins (Moon et al. 2016). Exercise was found to increase plasma CTSB levels in monkeys and humans (Moon et al. 2016), but a 20-week exercise intervention in children did not find any significant connection between CTSB and brain health outcomes (Rodriguez-Ayllon et al. 2023). Additionally, CTSB has been connected to brain-related functions in several mice studies (Sharanova et al. 2016; Z. Wang et al. 2018; Zhanaeva et al. 2018). For example, a mouse model for chronic social stress revealed increased activity of cathepsin В in the hypothalamus and nucleus caudatus with depressive-like behaviour (Zhanaeva et al. 2018). Contrarily, decreased cathepsin B activity was found after acute emotional stress in mice (Sharanova et al. 2016). CTSB shows a potential mediator role in the brain induced by physical and mental stressors which should be further investigated.

Other proteins significantly associated with the p-factor and directly connected to EGFR in our study were golgi membrane protein 1 (GOLM1) and S100 calcium binding protein A4 (S100A4). So far, only a few studies have described these proteins in relation to mental health. Increased *GOLM1* gene expression was found in soldiers with PTSD (Boscarino et al. 2019), while a recent study suggested that S1004A could be a protective factor against oxidative stress and brain injury mediating the antidepressant activity of ascorbic acid (vitamin C). This effect occurs through the activation of ErbB4-BDNF signalling pathway (Han et al. 2022; Pankratova et al. 2018). Particularly strong evidence supports the role of the Neuregulin-1 (NRG1)-ErbB4 signalling on synaptic plasticity (Ledonne and Mercuri 2019; Mei and Nave 2014). Neuregulins are a family of epidermal growth factor-related proteins acting on the ErbB tyrosine kinase receptors (Ledonne and Mercuri 2019; Mei and Nave 2014).

Associations of plasma reticulon-4 receptor-like 2 (RTN4L2), superoxide dismutase 2 (SOD2) and ficolin 3 (FCN3) with the p-factor were observed in this study. Reticulon-4 receptors (RTN4R), also known as NogoRs, are surface proteins expressed in neurons (J. Wang et al. 2021). RTN4Rs are involved in synaptogenesis and inhibition of axonal and dendrite growth, and thus neuronal plasticity (J. Wang et al. 2021; Willi and Schwab 2013). Human genetics studies have revealed the linkage between Nogo receptors and SCZ (Jitoku et al. 2011; Kimura et al. 2017; Willi and Schwab 2013). For example, a rare variant in *RTN4R*, affecting the formation of growth cones *in vitro,* was associated with SCZ (Kimura et al. 2017). The role of RTN4Rs in SCZ seems to be mediated by neurodevelopmental and myelin-related abnormalities (Willi and Schwab 2013). However, further studies are needed to clarify the exact role of RTN4Rs in mental health. SOD2 was found to play a role in neurodegenerative disease according to the Disease Ontology database, a polymorphism in the *sod2* gene was associated with differences in white matter microstructure and suboptimal brain aging (Salminen et al. 2017). Interestingly, ficolin activation was negatively associated with severity of SCZ (Gracia et al. 2021) and in our recent study, the plasma abundance of ficolin 2, a similar protein, was found to be positively correlated with the Strength and Difficulties Questionnaire (SDQ) score in adolescents.

Half of the proteins found to be significant in this study were connected to the extracellular matrix. HSPG2, FBLN1 and LAMB1 were also strongly connected to each other according to the STRING database, being structural components of the basal membrane, specifically in the brain. Coupled with proteins related to neuronal plasticity, proteins identified in this study may potentially relate to the previously noted inverse relationship between the p-factor and the microstructural integrity of white matter as observed through neuroimaging (Romer et al. 2021). Further studies are needed to investigate the possible connections of found proteins with the brain microstructure and functioning.

Large-scale proteomic studies with plasma samples can present multiple challenges that need to be addressed to generate robust and meaningful results. For instance, protein expression in plasma is dynamic and both interindividual and sample variability can be notable. Furthermore, plasma proteomic studies differ by the pipelines and methods used due to a lack of standard protocols (Ignjatovic et al. 2019). Additional challenges include ensuring consistent sample handling and processing (Bell et al. 2009), normalizing data, correcting signal drift and batch effects (Čuklina et al. 2021; Lazar et al. 2016), accounting for biological variability (Paul et al. 2013), improving reproducibility (Ioannidis 2011), and managing the resource-intensive nature of such studies (Nesvizhskii 2014). Despite these limitations, proteomics remains a powerful tool that can contribute to better diagnostics of mental health (Davalieva, Kostovska, and Dwork 2016; Rodrigues-Amorim et al. 2019). The major constraint in this study is that the proteomic data was only obtained once for each participant. This one-time snapshot of a dynamically evolving organism makes it challenging to conclusively link the identified biomarkers to the investigated p-factor. The true nature of these associations is hard to determine based solely on these data. These correlations could be the outcome of underlying biological processes or inherent biological traits of the participants, which might simultaneously influence both protein abundance and the p-factor (the observed behaviour). Alternatively, the changes in protein abundance and the p-factor could be causally related, either as a cause or as an effect. Furthermore, as demonstrated in the study by Shin and colleagues (Shin et al. 2021), mental conditions which contribute to distinct subcomponents of the p-factor, may cause divergent effects on the abundance of plasma proteins, suggesting that a more detailed investigation into the various components of the p-factor may be needed to identify more specific biomarkers.

The strength of this study lies in its large cohort size, and the use of modern proteomics methods, which made it possible to obtain proteome profiles of hundreds of individuals, each comprising hundreds of plasma protein abundancies. This large scale allows us to identify common patterns in the proteomes of individuals with high and low p-factor values. While the changes of plasma abundancies of some of the proteins were previously reported, other proteins were linked to a vulnerability to the development of general psychopathology for the first time. Our research utilized the FT12 cohort, a large and thoroughly characterized population-based cohort with a broad range of measured characteristics making the proteomic data gathered in this study an invaluable resource for future exploration and analysis.

## Conclusions

The study suggests that examining plasma proteomic profiles makes it possible to elucidate the biological processes related to the p-factor, which may inform the future development of novel screening, diagnostic or therapeutic strategies for mental disorders. The results revealed proteins with common cellular functions connected to the p-factor reflecting the general psychopathology. However, further studies are needed to examine the identified proteins and their potential as biomarkers for mental health dysfunction. In the future, utilization of the p-factor may also have implications for the development of interventions targeting common underlying factors that contribute to multiple forms of mental disorders. By addressing these shared factors, interventions could potentially be more effective in improving mental health outcomes across a range of disorders.

## Reporting summary

## Acknowledgements

We gratefully acknowledge the contribution of the Early Environmental quality and life-course mental health effects (Equal-Life) project team, the Turku Proteomics Facility team supported by Biocenter Finland for mass spectrometry, and the laboratory assistance of Ms. Mirka Tikkanen. We also thank Teemu Palviainen and Mia Urjansson for data and sample management, as well as Marja Heinonen-Guzejev and Gabin Drouard for their feedback on the manuscript.

## Author Contributions

KMK, ISM, JK and IvK designed the study. JK, AW LP and RJR guided the process and provided the samples. ISM, MI and AA pre-processed the samples. AMA performed statistical and bioinformatics analyses. A-KP, AMA, ISM and MI participated in planning and literature analyses. AMA, A-KP, AW and KMK drafted the manuscript. AW, JK, LP and RJR provided critical input to the draft manuscript. All authors read and approved the final version of the manuscript.

## Corresponding author

Correspondence and requests for materials should be addressed to Katja M. Kanninen.

## Additional information

Supplementary Information is available for this paper.

## Data availability

The data analyzed in this study is subject to the following licenses/restrictions: The FT12 data is not publicly available due to the restrictions of informed consent. Requests to access these datasets should be directed to the Institute for Molecular Medicine Finland (FIMM) Data Access Committee (DAC) for authorized researchers who have IRB/ethics approval and an institutionally approved study plan. To ensure the protection of privacy and compliance with national data protection legislation, a data use/transfer agreement is needed, the content and specific clauses of which will depend on the nature of the requested data.

## Supplementary data

**Supplementary figure 1.**
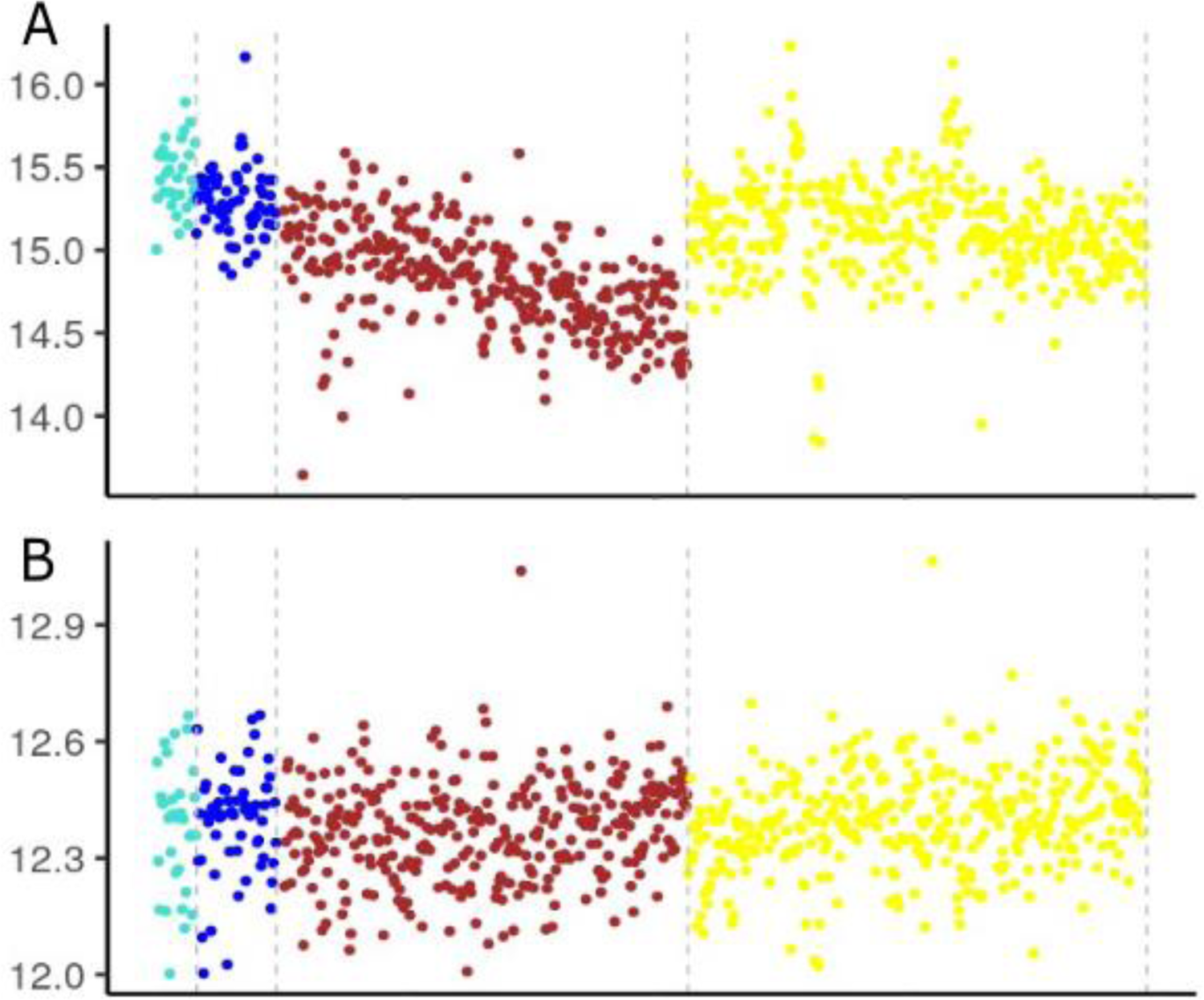
Plots of mean intensity values for each sample. A) the original values B) after drift and batch correction. Each dot represents the mean intensity values for single samples and the samples are plotted chronologically. The batches are colored and separated with dotted grey lines. The first two batches coincide with separate launches of the instrument. The latter batches are from the same launch, in-between the batches the instrument was cleaned, which led to a noticeable increase in the mean intensity.

**Supplementary figure 2.**
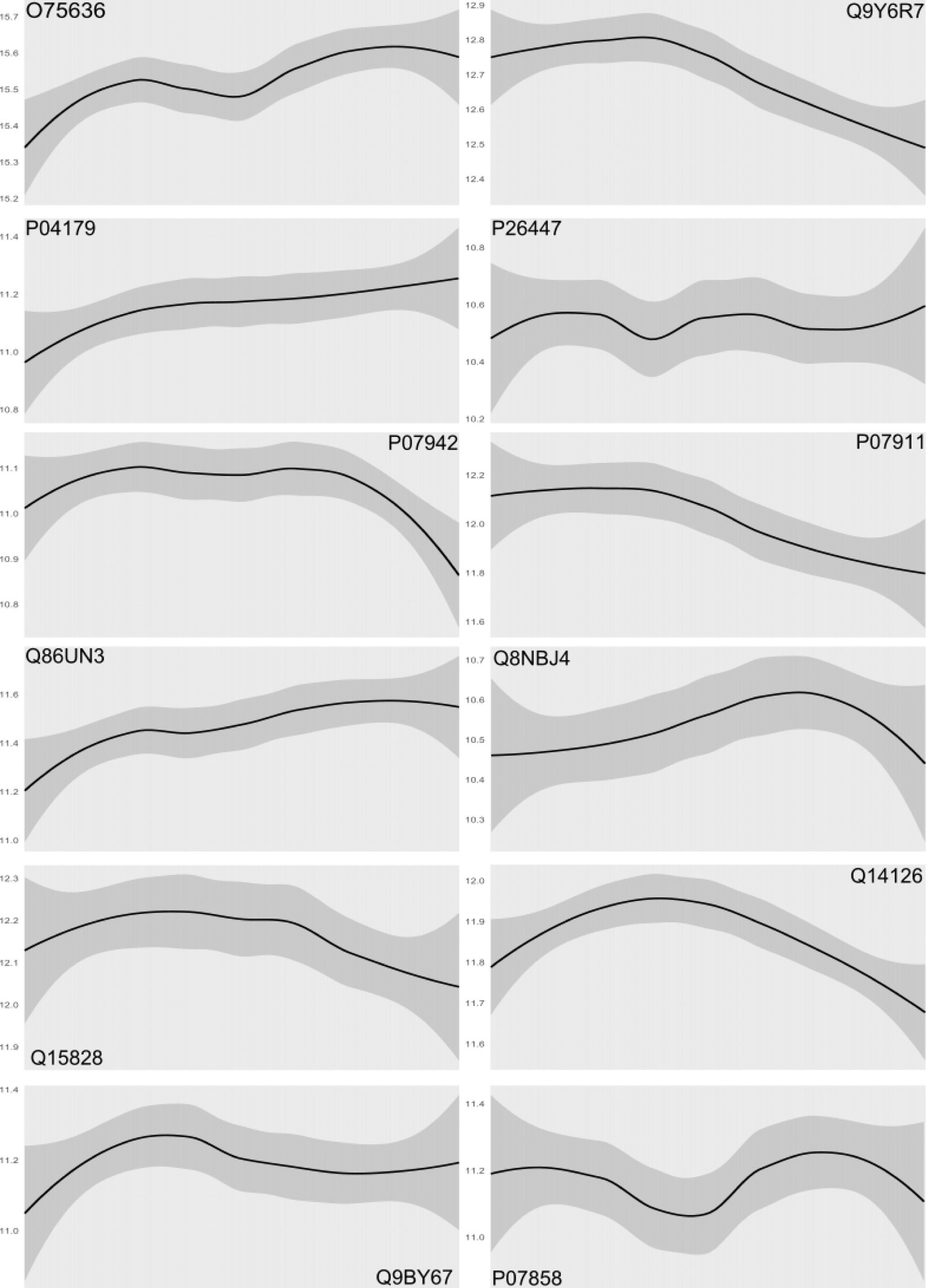
Plots of abundance for proteins with significant non-linear associations with the p-factor. The lines were constructed using geom_smooth function of the ggplot2 package. The abundance scales are different for each plot. X-axis represent participants sorted by the p-factor, lowest p-factor being on the left.

---

**Supplementary table 1.** The results of the enrichment analysis using STRINGdb

| *Category* | *Term* | *Protein number* | *Genes in background* | *Protein IDs* | *p-value* | *FDR* | *Description* |
| --- | --- | --- | --- | --- | --- | --- | --- |
| COMPARTMENTS | GOCC:0005576 | 11 | 2035 | LAMB1,FCN3,CST6,FBLN1,RTN4RL2,CTSB,S100A4,HSPG2, GOLM1,UMOD,SOD2 | < 0.001 | < 0.001 | Extracellular region |
| COMPARTMENTS | GOCC:0005615 | 8 | 985 | LAMB,FCN3,FBLN1,CTSB,S100A4,HSPG2,UMOD,SOD2 | < 0.001 | < 0.001 | Extracellular space |
| COMPARTMENTS | GOCC:0070062 | 5 | 368 | CTSB,S100A4,HSPG2,UMOD,SOD2 | < 0.001 | 0.003 | Extracellular exosome |
| COMPARTMENTS | GOCC:0031012 | 4 | 253 | LAMB1,FBLN1,S100A4,HSPG2 | < 0.001 | 0.010 | Extracellular matrix |
| Component | GO:0005615 | 13 | 3195 | LAMB1,DSG2,FCN3,CST6,FBLN1,RTN4RL2,CTSB,S100A4,HSPG2,GOLM1,UMOD,SOD2,FCGBP | < 0.001 | < 0.001 | Extracellular space |
| Component | GO:0070062 | 11 | 2099 | LAMB1,DSG2,CST6,FBLN1,RTN4RL2,CTSB,S100A4,HSPG2, UMOD,SOD2,FCGBP | < 0.001 | < 0.001 | Extracellular exosome |
| Component | GO:0031012 | 7 | 527 | LAMB1,FCN3,FBLN1,RTN4RL2,CTSB,S100A4,HSPG2 | < 0.001 | < 0.001 | Extracellular matrix |
| Component | GO:0062023 | 6 | 396 | LAMB1,FCN3,FBLN1,CTSB,S100A4,HSPG2 | < 0.001 | < 0.001 | Collagen-containing extracellular matrix |
| Component | GO:0005604 | 3 | 96 | LAMB1,FBLN1,HSPG2 | < 0.001 | 0.008 | Basement membrane |
| Function | GO:0005201 | 4 | 119 | LAMB1,FBLN1,HSPG2,UMOD | < 0.001 | 0.005 | Extracellular matrix structural constituent |
| TISSUES | BTO:0001491 | 12 | 5020 | LAMB1,DSG2,FCN3,CADM1,FBLN1,CTSB,S100A4,HSPG2, GOLM1,UMOD,SOD2,FCGBP | < 0.001 | 0.008 | Viscus |
| TISSUES | BTO:0004850 | 4 | 170 | LAMB1,DSG2,HSPG2,FCGBP | < 0.001 | 0.008 | Bone marrow cell |
| Keyword | KW-0732 | 11 | 3233 | LAMB1,DSG2,FCN3,CST6,CADM1,FBLN1,RTN4RL2,CTSB,HSPG2,UMOD,FCGBP | < 0.001 | < 0.001 | Signal |
| Keyword | KW-0325 | 11 | 4349 | LAMB1,DSG2,FCN3,CST6,CADM1,FBLN1,RTN4RL2,CTSB,HSPG2,GOLM1,UMOD | < 0.001 | 0.004 | Glycoprotein |
| Keyword | KW-0964 | 7 | 1818 | LAMB1,FCN3,CST6,FBLN1,CTSB,HSPG2,UMOD | < 0.001 | 0.022 | Secreted |
| Keyword | KW-1015 | 9 | 3304 | LAMB1,FCN3,CST6,CADM1,FBLN1,RTN4RL2,CTSB,HSPG2,UMOD | < 0.001 | 0.022 | Disulfide bond |
| Keyword | KW-0106 | 5 | 875 | DSG2,FCN3,FBLN1,S100A4,HSPG2 | < 0.001 | 0.031 | Calcium |
| Keyword | KW-0424 | 2 | 30 | LAMB1,HSPG2 | < 0.001 | 0.031 | Laminin EGF-like domain |
| Keyword | KW-0084 | 2 | 39 | LAMB1,HSPG2 | < 0.001 | 0.037 | Basement membrane |
| Keyword | KW-0245 | 3 | 229 | FBLN1,HSPG2,UMOD | 0.001 | 0.046 | EGF-like domain |
| InterPro | IPR000742 | 5 | 233 | LAMB1,FBLN1,HSPG2,UMOD,FCGBP | < 0.001 | < 0.001 | EGF-like domain |

## References

Bateman, Alex, Maria-Jesus Martin, Sandra Orchard, Michele Magrane, Shadab Ahmad, Emanuele Alpi, Emily H Bowler-Barnett, et al. 2023. “UniProt: The Universal Protein Knowledgebase in 2023.” Nucleic Acids Research 51 (D1): D523–31. https://doi.org/10.1093/nar/gkac1052.

Bell, Alexander W, Eric W Deutsch, Catherine E Au, Robert E Kearney, Ron Beavis, Salvatore Sechi, Tommy Nilsson, and John J M Bergeron. 2009. “A HUPO Test Sample Study Reveals Common Problems in Mass Spectrometry–Based Proteomics.” Nature Methods 6 (6): 423–30. https://doi.org/10.1038/nmeth.1333.

Benjamini, Yoav, and Yosef Hochberg. 1995. “Controlling the False Discovery Rate: A Practical and Powerful Approach to Multiple Testing.” Journal of the Royal Statistical Society: Series B (Methodological) 57 (1): 289–300. https://doi.org/10.1111/j.2517-6161.1995.tb02031.x.

Birchmeier, Carmen. 2009. “ErbB Receptors and the Development of the Nervous System.” Experimental Cell Research 315 (4): 611–18. https://doi.org/10.1016/j.yexcr.2008.10.035.

Bishop, Joseph R., Manuela Schuksz, and Jeffrey D. Esko. 2007. “Heparan Sulphate Proteoglycans Fine-Tune Mammalian Physiology.” Nature 446 (7139): 1030–37. https://doi.org/10.1038/nature05817.

Bohlega, Saeed, Huda Al-Ajlan, and Amr Al-Saif. 2014. “Mutation of Fibulin-1 Causes a Novel Syndrome Involving the Central Nervous System and Connective Tissues.” European Journal of Human Genetics 22 (5): 640–43. https://doi.org/10.1038/ejhg.2013.210.

Borges, Jazlyn P., Katrina Mekhail, Gregory D. Fairn, Costin N. Antonescu, and Benjamin E. Steinberg. 2021. “Modulation of Pathological Pain by Epidermal Growth Factor Receptor.” Frontiers in Pharmacology 12 (May). https://doi.org/10.3389/fphar.2021.642820.

Boscarino, Cathy, Thomas Nalpathamkalam, Giovanna Pellecchia, Weili Li, Bhooma Thiruvahindrapuram, and Daniele Merico. 2019. “Using Next-Generation Sequencing Transcriptomics To Determine Markers of Post-Traumatic Symptoms: Preliminary Findings from a Post-Deployment Cohort of Soldiers.” G3 Genes|Genomes|Genetics 9 (2): 463–71. https://doi.org/10.1534/g3.118.200516.

Campeau, Anaamika, Robert H. Mills, Toer Stevens, Leigh-Ana Rossitto, Michael Meehan, Pieter Dorrestein, Rebecca Daly, et al. 2022. “Multi-Omics of Human Plasma Reveals Molecular Features of Dysregulated Inflammation and Accelerated Aging in Schizophrenia.” Molecular Psychiatry 27 (2): 1217–25. https://doi.org/10.1038/s41380-021-01339-z.

Caspi, Avshalom, Renate M. Houts, Daniel W. Belsky, Sidra J. Goldman-Mellor, HonaLee Harrington, Salomon Israel, Madeline H. Meier, et al. 2014. “The p Factor: One General Psychopathology Factor in the Structure of Psychiatric Disorders?” Clinical Psychological Science 2 (2): 119–37. https://doi.org/10.1177/2167702613497473.

Chen, Yi-Hua, Neng-Yuan Hu, Ding-Yu Wu, Lin-Lin Bi, Zheng-Yi Luo, Lang Huang, Jian-Lin Wu, et al. 2022. “PV Network Plasticity Mediated by Neuregulin1-ErbB4 Signalling Controls Fear Extinction.” Molecular Psychiatry 27 (2): 896–906. https://doi.org/10.1038/s41380-021-01355-z.

Comes, Ashley L., Sergi Papiol, Thorsten Mueller, Philipp E. Geyer, Matthias Mann, and Thomas G. Schulze. 2018. “Proteomics for Blood Biomarker Exploration of Severe Mental Illness: Pitfalls of the Past and Potential for the Future.” Translational Psychiatry 8 (1): 160. https://doi.org/10.1038/S41398-018-0219-2.

Cooley, Marion A., Christine B. Kern, Victor M. Fresco, Andy Wessels, Robert P. Thompson, Tim C. McQuinn, Waleed O. Twal, Corey H. Mjaatvedt, Christopher J. Drake, and W. Scott Argraves. 2008. “Fibulin-1 Is Required for Morphogenesis of Neural Crest-Derived Structures.” Developmental Biology 319 (2): 336–45. https://doi.org/10.1016/j.ydbio.2008.04.029.

Čuklina, Jelena, Chloe H Lee, Evan G Williams, Tatjana Sajic, Ben C Collins, María Rodríguez Martínez, Varun S Sharma, et al. 2021. “Diagnostics and Correction of Batch Effects in Large-scale Proteomic Studies: A Tutorial.” Molecular Systems Biology 17 (8). https://doi.org/10.15252/msb.202110240.

Davalieva, Katarina, Ivana Maleva Kostovska, and Andrew J. Dwork. 2016. “Proteomics Research in Schizophrenia.” Frontiers in Cellular Neuroscience 10 (February). https://doi.org/10.3389/FNCEL.2016.00018.

Domenici, Enrico, David R. Willé, Federica Tozzi, Inga Prokopenko, Sam Miller, Astrid McKeown, Claire Brittain, et al. 2010. “Plasma Protein Biomarkers for Depression and Schizophrenia by Multi Analyte Profiling of Case-Control Collections.” Edited by Katharina Domschke. PLoS ONE 5 (2): e9166. https://doi.org/10.1371/journal.pone.0009166.

Fernandes, Brisa S., Yulin Dai, Peilin Jia, and Zhongming Zhao. 2022. “Charting the Proteome Landscape in Major Psychiatric Disorders: From Biomarkers to Biological Pathways towards Drug Discovery.” European Neuropsychopharmacology 61 (August): 43–59. https://doi.org/10.1016/j.euroneuro.2022.06.001.

Fiori, Laura M., Aron Kos, Rixing Lin, Jean-Francois Théroux, Juan Pablo Lopez, Claudia Kühne, Carola Eggert, et al. 2021. “MiR-323a Regulates ERBB4 and Is Involved in Depression.” Molecular Psychiatry 26 (8): 4191–4204. https://doi.org/10.1038/s41380-020-00953-7.

Frankenfield, Ashley M., Jiawei Ni, Mustafa Ahmed, and Ling Hao. 2022. “Protein Contaminants Matter: Building Universal Protein Contaminant Libraries for DDA and DIA Proteomics.” Journal of Proteome Research 21 (9): 2104–13. https://doi.org/10.1021/acs.jproteome.2c00145.

García-Gutiérrez, Maria Salud, Francisco Navarrete, Francisco Sala, Ani Gasparyan, Amaya Austrich-Olivares, and Jorge Manzanares. 2020. “Biomarkers in Psychiatry: Concept, Definition, Types and Relevance to the Clinical Reality.” Frontiers in Psychiatry 11 (May). https://doi.org/10.3389/FPSYT.2020.00432.

Gracia Diego Fabian Karvat, Eloisa Maria Pontarolo Gomes, Tamires Amelotti Coelho, Marcelo Carriello, Fabiana Antunes de Andrade, Lorena Bavia, Iara Jose Messias-Reason, and Raffael Massuda. 2021. “Ficolin Activation as a Potential Biomarker of the Severity of Schizophrenia.” Psychiatry Research 304 (October): 114122. https://doi.org/10.1016/j.psychres.2021.114122.

Han, Qian-Qian, Peng-Fei Wu, Yi-Heng Li, Yu Cao, Jian-Guo Chen, and Fang Wang. 2022. “SVCT2–Mediated Ascorbic Acid Uptake Buffers Stress Responses via DNA Hydroxymethylation Reprogramming of S100 Calcium-Binding Protein A4 Gene.” Redox Biology 58 (December): 102543. https://doi.org/10.1016/j.redox.2022.102543.

Hartlieb, Eva, Vera Rötzer, Mariya Radeva, Volker Spindler, and Jens Waschke. 2014. “Desmoglein 2 Compensates for Desmoglein 3 but Does Not Control Cell Adhesion via Regulation of P38 Mitogen-Activated Protein Kinase in Keratinocytes.” Journal of Biological Chemistry 289 (24): 17043–53. https://doi.org/10.1074/jbc.M113.489336.

Hartwig, Fernando Pires, Maria Carolina Borges, Bernardo Lessa Horta, Jack Bowden, and George Davey Smith. 2017. “Inflammatory Biomarkers and Risk of Schizophrenia: A 2-Sample Mendelian Randomization Study.” JAMA Psychiatry 74 (12): 1226–33. https://doi.org/10.1001/JAMAPSYCHIATRY.2017.3191.

Haywood, Darren, Frank D. Baughman, Barbara A. Mullan, and Karen R. Heslop. 2022. “What Accounts for the Factors of Psychopathology? An Investigation of the Neurocognitive Correlates of Internalising, Externalising, and the p-Factor.” Brain Sciences 12 (4): 421. https://doi.org/10.3390/brainsci12040421.

*Health at a Glance: Europe* 2022. 2022. Health at a Glance: Europe. OECD. https://doi.org/10.1787/507433b0-en.

“Human Protein Atlas.” 2022. 2022. proteinatlas.org.

Ignjatovic, Vera, Philipp E. Geyer, Krishnan K. Palaniappan, Jessica E. Chaaban, Gilbert S. Omenn, Mark S. Baker, Eric W. Deutsch, and Jochen M. Schwenk. 2019. “Mass Spectrometry-Based Plasma Proteomics: Considerations from Sample Collection to Achieving Translational Data.” Journal of Proteome Research 18 (12): 4085–97. https://doi.org/10.1021/acs.jproteome.9b00503.

Ioannidis, John P. A. 2011. “Comparison of Effect Sizes Associated With Biomarkers Reported in Highly Cited Individual Articles and in Subsequent Meta-Analyses.” JAMA 305 (21): 2200. https://doi.org/10.1001/jama.2011.713.

Jensen, Arthur R. 1999. “The g Factor: The Science of Mental Ability.” Psycoloquy 10 (04): 36–2443.

Ji, Ellen, Danny Boerrigter, Helen Q. Cai, David Lloyd, Jason Bruggemann, Maryanne O’Donnell, Cherrie Galletly, et al. 2022. “Peripheral Complement Is Increased in Schizophrenia and Inversely Related to Cortical Thickness.” Brain, Behavior, and Immunity 101 (March): 423–34. https://doi.org/10.1016/j.bbi.2021.11.014.

Jitoku, Daisuke, Eiji Hattori, Yoshimi Iwayama, Kazuo Yamada, Tomoko Toyota, Mitsuru Kikuchi, Motoko Maekawa, Toru Nishikawa, and Takeo Yoshikawa. 2011. “Association Study of Nogo-Related Genes with Schizophrenia in a Japanese Case-Control Sample.” American Journal of Medical Genetics Part B: Neuropsychiatric Genetics 156 (5): 581–92. https://doi.org/10.1002/ajmg.b.31199.

Kaprio, Jaakko. 2006. “Twin Studies in Finland 2006.” Twin Research and Human Genetics 9 (6): 772–77. https://doi.org/10.1375/183242706779462778.

Kim, Young Jong, Jin Kyung Park, Won Sub Kang, Su Kang Kim, Hae Jeong Park, Min Nam, and Jong Woo Kim. 2015. “LAMB1 Polymorphism Is Associated with Autism Symptom Severity in Korean Autism Spectrum Disorder Patients.” Nordic Journal of Psychiatry 69 (8): 594–98. https://doi.org/10.3109/08039488.2015.1022597.

Kimura, H, Y Fujita, T Kawabata, K Ishizuka, C Wang, Y Iwayama, Y Okahisa, et al. 2017. “A Novel Rare Variant R292H in RTN4R Affects Growth Cone Formation and Possibly Contributes to Schizophrenia Susceptibility.” Translational Psychiatry 7 (8): e1214–e1214. https://doi.org/10.1038/tp.2017.170.

Laceulle, Odilia M., Wilma A. M. Vollebergh, and Johan Ormel. 2015. “The Structure of Psychopathology in Adolescence.” Clinical Psychological Science 3 (6): 850–60. https://doi.org/10.1177/2167702614560750.

Lahey, Benjamin B., Brooks Applegate, Jahn K. Hakes, David H. Zald, Ahmad R. Hariri, and Paul J. Rathouz. 2012. “Is There a General Factor of Prevalent Psychopathology during Adulthood?” Journal of Abnormal Psychology 121 (4): 971–77. https://doi.org/10.1037/a0028355.

Lazar, Cosmin, Laurent Gatto, Myriam Ferro, Christophe Bruley, and Thomas Burger. 2016. “Accounting for the Multiple Natures of Missing Values in Label-Free Quantitative Proteomics Data Sets to Compare Imputation Strategies.” Journal of Proteome Research 15 (4): 1116–25. https://doi.org/10.1021/acs.jproteome.5b00981.

Ledonne, Ada, and Nicola B. Mercuri. 2019. “On the Modulatory Roles of Neuregulins/ErbB Signaling on Synaptic Plasticity.” International Journal of Molecular Sciences 21 (1): 275. https://doi.org/10.3390/ijms21010275.

Li, Zhen-Zhen, Wen-Juan Han, Zhi-Chuan Sun, Yun Chen, Jun-Yi Sun, Guo-Hong Cai, Wan-Neng Liu, et al. 2021. “Extracellular Matrix Protein Laminin Β1 Regulates Pain Sensitivity and Anxiodepression-like Behaviors in Mice.” Journal of Clinical Investigation 131 (15). https://doi.org/10.1172/JCI146323.

Liu, Mingyi, and Ashok Dongre. 2021. “Proper Imputation of Missing Values in Proteomics Datasets for Differential Expression Analysis.” Briefings in Bioinformatics 22 (3). https://doi.org/10.1093/BIB/BBAA112.

Long, Qing, Rui Wang, Maoyang Feng, Xinling Zhao, Yilin Liu, Xiao Ma, Lei Yu, et al. 2021. “Construction and Analysis of a Diagnostic Model Based on Differential Expression Genes in Patients With Major Depressive Disorder.” Frontiers in Psychiatry 12 (December). https://doi.org/10.3389/fpsyt.2021.762683.

MacNeil, Raymond R., and Daniel J. Müller. 2016. “Genetics of Common Antipsychotic-Induced Adverse Effects.” Complex Psychiatry 2 (2): 61–78. https://doi.org/10.1159/000445802.

Malik, Sahil, Ravinder Singh, Govind Arora, Akriti Dangol, and Sanjay Goyal. 2021. “Biomarkers of Major Depressive Disorder: Knowing Is Half the Battle.” Clinical Psychopharmacology and Neuroscience 19 (1): 12–25. https://doi.org/10.9758/cpn.2021.19.1.12.

Mei, Lin, and Klaus-Armin Nave. 2014. “Neuregulin-ERBB Signaling in the Nervous System and Neuropsychiatric Diseases.” Neuron 83 (1): 27–49. https://doi.org/10.1016/j.neuron.2014.06.007.

Mei, Lin, and Wen-Cheng Xiong. 2008. “Neuregulin 1 in Neural Development, Synaptic Plasticity and Schizophrenia.” Nature Reviews Neuroscience 9 (6): 437–52. https://doi.org/10.1038/nrn2392.

Moon, Hyo Youl, Andreas Becke, David Berron, Benjamin Becker, Nirnath Sah, Galit Benoni, Emma Janke, et al. 2016. “Running-Induced Systemic Cathepsin B Secretion Is Associated with Memory Function.” Cell Metabolism 24 (2): 332–40. https://doi.org/10.1016/j.cmet.2016.05.025.

Munn-Chernoff, Melissa A., Emma C. Johnson, Yi-Ling Chou, Jonathan R.I. Coleman, Laura M. Thornton, Raymond K. Walters, Zeynep Yilmaz, et al. 2021. “Shared Genetic Risk between Eating Disorder- and Substance-use-related Phenotypes: Evidence from Genome-wide Association Studies.” Addiction Biology 26 (1). https://doi.org/10.1111/adb.12880.

Nakamura, Kuniyuki, Tomoko Ikeuchi, Kazuki Nara, Craig S. Rhodes, Peipei Zhang, Yuta Chiba, Saiko Kazuno, et al. 2019. “Perlecan Regulates Pericyte Dynamics in the Maintenance and Repair of the Blood–Brain Barrier.” Journal of Cell Biology 218 (10): 3506–25. https://doi.org/10.1083/jcb.201807178.

Nawwar, Dalia A., Hala F. Zaki, and Rabab H. Sayed. 2022. “Role of the NRG1/ErbB4 and PI3K/AKT/MTOR Signaling Pathways in the Anti-Psychotic Effects of Aripiprazole and Sertindole in Ketamine-Induced Schizophrenia-like Behaviors in Rats.” Inflammopharmacology 30 (5): 1891–1907. https://doi.org/10.1007/s10787-022-01031-w.

Nesvizhskii, Alexey I. 2014. “Proteogenomics: Concepts, Applications and Computational Strategies.” Nature Methods 11 (11): 1114–25. https://doi.org/10.1038/nmeth.3144.

Ohsawa, Ikuroh, Chizuko Takamura, and Shinichi Kohsaka. 2001. “Fibulin-1 Binds the Amino-Terminal Head of β-Amyloid Precursor Protein and Modulates Its Physiological Function.” Journal of Neurochemistry 76 (5): 1411–20. https://doi.org/10.1046/j.1471-4159.2001.00144.x.

Oltmanns, Joshua R., Gregory T. Smith, Thomas F. Oltmanns, and Thomas A. Widiger. 2018. “General Factors of Psychopathology, Personality, and Personality Disorder: Across Domain Comparisons.” Clinical Psychological Science 6 (4): 581–89. https://doi.org/10.1177/2167702617750150.

Organization, World Health. 2022. “World Mental Health Report: Transforming Mental Health for All.”

Orlando, Eleonora, and Ruedi Aebersold. 2019. “On the Contribution of Mass Spectrometry-Based Platforms to the Field of Personalized Oncology.” TrAC Trends in Analytical Chemistry 110 (January): 129–42. https://doi.org/10.1016/j.trac.2018.10.018.

Pankratova, Stanislava, Jorg Klingelhofer, Oksana Dmytriyeva, Sylwia Owczarek, Alexander Renziehausen, Nelofer Syed, Alexandra E. Porter, David T. Dexter, and Darya Kiryushko. 2018. “The S100A4 Protein Signals through the ErbB4 Receptor to Promote Neuronal Survival.” Theranostics 8 (14): 3977–90. https://doi.org/10.7150/thno.22274.

Paul, Debasish, Avinash Kumar, Akshada Gajbhiye, Manas K. Santra, and Rapole Srikanth. 2013. “Mass Spectrometry-Based Proteomics in Molecular Diagnostics: Discovery of Cancer Biomarkers Using Tissue Culture.” BioMed Research International 2013: 1–16. https://doi.org/10.1155/2013/783131.

Pham, Thang V, Alex A Henneman, and Connie R Jimenez. 2020. “Iq: An R Package to Estimate Relative Protein Abundances from Ion Quantification in DIA-MS-Based Proteomics.” Edited by Alfonso Valencia. Bioinformatics 36 (8): 2611–13. https://doi.org/10.1093/bioinformatics/btz961.

Phipson, Belinda, Stanley Lee, Ian J. Majewski, Warren S. Alexander, and Gordon K. Smyth. 2016. “Robust Hyperparameter Estimation Protects against Hypervariable Genes and Improves Power to Detect Differential Expression.” The Annals of Applied Statistics 10 (2). https://doi.org/10.1214/16-AOAS920.

Pulkkinen, Lea. 2017. Human Development from Middle Childhood to Middle Adulthood. London: Routledge. https://doi.org/10.4324/9781315732947.

Pulkkinen, Lea, Jaakko Kaprio, and Richard J Rose. 1999. “Peers, Teachers and Parents as Assessors of the Behavioural and Emotional Problems of Twins and Their Adjustment: The Multidimensional Peer Nomination Inventory.” Twin Research (1999) 2 (4): 274–85. https://doi.org/10.1375/136905299320565762.

Ritchie, Matthew E., Belinda Phipson, Di Wu, Yifang Hu, Charity W. Law, Wei Shi, and Gordon K. Smyth. 2015. “Limma Powers Differential Expression Analyses for RNA-Sequencing and Microarray Studies.” Nucleic Acids Research 43 (7): e47–e47. https://doi.org/10.1093/nar/gkv007.

Rodrigues-Amorim, Daniela, Tania Rivera-Baltanás, María del Carmen Vallejo-Curto, Cynthia Rodriguez-Jamardo, Elena de las Heras, Carolina Barreiro-Villar, María Blanco-Formoso, et al. 2019. “Proteomics in Schizophrenia: A Gateway to Discover Potential Biomarkers of Psychoneuroimmune Pathways.” Frontiers in Psychiatry 10 (November). https://doi.org/10.3389/FPSYT.2019.00885.

Rodriguez-Ayllon, María, Abel Plaza-Florido, Andrea Mendez-Gutierrez, Signe Altmäe, Patricio Solis-Urra, Concepción M. Aguilera, Andrés Catena, Francisco B. Ortega, and Irene Esteban-Cornejo. 2023. “The Effects of a 20-Week Exercise Program on Blood-Circulating Biomarkers Related to Brain Health in Overweight or Obese Children: The ActiveBrains Project.” Journal of Sport and Health Science 12 (2): 175–85. https://doi.org/10.1016/j.jshs.2022.12.007.

Rodríguez Cerdeira, Carmen, Elena Sánchez-Blanco, Beatriz Sánchez-Blanco, and Jose Luis González-Cespón. 2017. “Protein Biomarkers of Mood Disorders.” International Journal of Immunopathology and Pharmacology 30 (1): 7–12. https://doi.org/10.1177/0394632016681017.

Romer, Adrienne L., Annchen R. Knodt, Maria L. Sison, David Ireland, Renate Houts, Sandhya Ramrakha, Richie Poulton, et al. 2021. “Replicability of Structural Brain Alterations Associated with General Psychopathology: Evidence from a Population-Representative Birth Cohort.” Molecular Psychiatry 26 (8): 3839–46. https://doi.org/10.1038/s41380-019-0621-z.

Ronald, Angelica. 2019. “Editorial: The Psychopathology p Factor: Will It Revolutionise the Science and Practice of Child and Adolescent Psychiatry?” Journal of Child Psychology and Psychiatry 60 (5): 497–99. https://doi.org/10.1111/jcpp.13063.

Rose, Richard J., Jessica E. Salvatore, Sari Aaltonen, Peter B. Barr, Leonie H. Bogl, Holly A. Byers, Kauko Heikkilä, et al. 2019. “FinnTwin12 Cohort: An Updated Review.” Twin Research and Human Genetics 22 (5): 302–11. https://doi.org/10.1017/thg.2019.83.

Salminen, Lauren E., Peter R. Schofield, Kerrie D. Pierce, Steven E. Bruce, Michael G. Griffin, David F. Tate, Ryan P. Cabeen, et al. 2017. “Vulnerability of White Matter Tracts and Cognition to the SOD2 Polymorphism: A Preliminary Study of Antioxidant Defense Genes in Brain Aging.” Behavioural Brain Research 329 (June): 111–19. https://doi.org/10.1016/j.bbr.2017.04.041.

Schriml, Lynn M, James B Munro, Mike Schor, Dustin Olley, Carrie McCracken, Victor Felix, J Allen Baron, et al. 2022. “The Human Disease Ontology 2022 Update.” Nucleic Acids Research 50 (D1): D1255–61. https://doi.org/10.1093/nar/gkab1063.

Shanmugan, Sheila, Daniel H. Wolf, Monica E. Calkins, Tyler M. Moore, Kosha Ruparel, Ryan D. Hopson, Simon N. Vandekar, et al. 2016. “Common and Dissociable Mechanisms of Executive System Dysfunction Across Psychiatric Disorders in Youth.” American Journal of Psychiatry 173 (5): 517–26. https://doi.org/10.1176/appi.ajp.2015.15060725.

Sharanova, N. E., N. V. Kirbaeva, I. Yu. Toropygin, E. V. Khryapova, E. V. Koplik, C. Kh. Soto, S. S. Pertsov, and A. V. Vasiliev. 2016. “Effect of Acute Emotional Stress on Proteomic Profile of Selected Brain Areas and Lysosomal Proteolysis in Rats with Different Behavioral Activity.” Bulletin of Experimental Biology and Medicine 161 (3): 355–58. https://doi.org/10.1007/s10517-016-3413-3.

Shi, Liang, and Clare M. Bergson. 2020. “Neuregulin 1: An Intriguing Therapeutic Target for Neurodevelopmental Disorders.” Translational Psychiatry 10 (1): 190. https://doi.org/10.1038/s41398-020-00868-5.

Shin, Dongyoon, Sang Jin Rhee, Jihyeon Lee, Injoon Yeo, Misol Do, Eun-Jeong Joo, Hee Yeon Jung, et al. 2021. “Quantitative Proteomic Approach for Discriminating Major Depressive Disorder and Bipolar Disorder by Multiple Reaction Monitoring-Mass Spectrometry.” Journal of Proteome Research 20 (6): 3188–3203. https://doi.org/10.1021/acs.jproteome.1c00058.

Smith, Gregory T., Emily A. Atkinson, Heather A. Davis, Elizabeth N. Riley, and Joshua R. Oltmanns. 2020. “The General Factor of Psychopathology.” Annual Review of Clinical Psychology 16 (1): 75–98. https://doi.org/10.1146/annurev-clinpsy-071119-115848.

Sun, Yuh-Man, Megan Cooper, Sophie Finch, Hsuan-Hwai Lin, Zhou-Feng Chen, Brenda P. Williams, and Noel J. Buckley. 2008. “Rest-Mediated Regulation of Extracellular Matrix Is Crucial for Neural Development.” Edited by Patrick Callaerts. PLoS ONE 3 (11): e3656. https://doi.org/10.1371/journal.pone.0003656.

Szklarczyk, Damian, Annika L Gable, David Lyon, Alexander Junge, Stefan Wyder, Jaime Huerta-Cepas, Milan Simonovic, et al. 2019. “STRING V11: Protein–Protein Association Networks with Increased Coverage, Supporting Functional Discovery in Genome-Wide Experimental Datasets.” Nucleic Acids Research 47 (D1): D607–13. https://doi.org/10.1093/nar/gky1131.

Szklarczyk, Damian, Annika L Gable, Katerina C Nastou, David Lyon, Rebecca Kirsch, Sampo Pyysalo, Nadezhda T Doncheva, et al. 2021. “The STRING Database in 2021: Customizable Protein–Protein Networks, and Functional Characterization of User-Uploaded Gene/Measurement Sets.” Nucleic Acids Research 49 (D1): D605–12. https://doi.org/10.1093/nar/gkaa1074.

Tordera, R.M., A.L. Garcia-García, N. Elizalde, V. Segura, E. Aso, E. Venzala, M.J. Ramírez, and J. Del Rio. 2011. “Chronic Stress and Impaired Glutamate Function Elicit a Depressive-like Phenotype and Common Changes in Gene Expression in the Mouse Frontal Cortex.” European Neuropsychopharmacology 21 (1): 23–32. https://doi.org/10.1016/j.euroneuro.2010.06.016.

Uhlén, Mathias, Max J. Karlsson, Andreas Hober, Anne-Sophie Svensson, Julia Scheffel, David Kotol, Wen Zhong, et al. 2019. “The Human Secretome.” Science Signaling 12 (609). https://doi.org/10.1126/scisignal.aaz0274.

Walters, Raymond K., Renato Polimanti, Emma C. Johnson, Jeanette N. McClintick, Mark J. Adams, Amy E. Adkins, Fazil Aliev, et al. 2018. “Transancestral GWAS of Alcohol Dependence Reveals Common Genetic Underpinnings with Psychiatric Disorders.” Nature Neuroscience 21 (12): 1656–69. https://doi.org/10.1038/s41593-018-0275-1.

Wang, Jie, Yi Miao, Rebecca Wicklein, Zijun Sun, Jinzhao Wang, Kevin M. Jude, Ricardo A. Fernandes, et al. 2021. “RTN4/NoGo-Receptor Binding to BAI Adhesion-GPCRs Regulates Neuronal Development.” Cell 184 (24): 5869–5885.e25. https://doi.org/10.1016/j.cell.2021.10.016.

Wang, Zhichao, Ping Li, Tong Wu, Shuangyue Zhu, Libin Deng, and Guangcheng Cui. 2018. “Axon Guidance Pathway Genes Are Associated with Schizophrenia Risk.” Experimental and Therapeutic Medicine 16 (6): 4519–26. https://doi.org/10.3892/ETM.2018.6781.

Whipp, Alyce M., Marja Heinonen-Guzejev, Kirsi H. Pietiläinen, Irene van Kamp, and Jaakko Kaprio. 2022. “Branched-Chain Amino Acids Linked to Depression in Young Adults.” Frontiers in Neuroscience 16 (September). https://doi.org/10.3389/fnins.2022.935858.

Wickham, Hadley. 2009. Ggplot2. Ggplot2. New York, NY: Springer New York. https://doi.org/10.1007/978-0-387-98141-3.

Willi, Roman, and Martin E. Schwab. 2013. “Nogo and Nogo Receptor: Relevance to Schizophrenia?” Neurobiology of Disease 54 (June): 150–57. https://doi.org/10.1016/j.nbd.2013.01.011.

World Health Organization. 2021. “Adolescent Mental Health.” November 17, 2021. https://www.who.int/news-room/fact-sheets/detail/adolescent-mental-health.

Yang, Ying C, Yun L Ma, Wen T Liu, and Eminy HY Lee. 2011. “Laminin-Β1 Impairs Spatial Learning through Inhibition of ERK/MAPK and SGK1 Signaling.” Neuropsychopharmacology 36 (12): 2571–86. https://doi.org/10.1038/npp.2011.148.

Zai, Clement C., Frankie H. Lee, Arun K. Tiwari, Justin Y. Lu, Vincenzo de Luca, Miriam S. Maes, Deanna Herbert, et al. 2018. “Investigation of the HSPG2 Gene in Tardive Dyskinesia – New Data and Meta-Analysis.” Frontiers in Pharmacology 9 (September). https://doi.org/10.3389/fphar.2018.00974.

Zai, Clement C., Miriam S. Maes, Arun K. Tiwari, Gwyneth C. Zai, Gary Remington, and James L. Kennedy. 2018. “Genetics of Tardive Dyskinesia: Promising Leads and Ways Forward.” Journal of the Neurological Sciences 389 (June): 28–34. https://doi.org/10.1016/j.jns.2018.02.011.

Zhanaeva, S. Ya., A. A. Rogozhnikova, E. L. Alperina, M. M. Gevorgyan, and G. V. Idov. 2018. “Changes in Activity of Cysteine Cathepsins B and L in Brain Structures of Mice with Aggressive and Depressive-Like Behavior Formed under Conditions of Social Stress.” Bulletin of Experimental Biology and Medicine 164 (4): 425–29. https://doi.org/10.1007/s10517-018-4004-2.

Zhou, Bin, Zhe Zhou, Yuling Chen, Haiteng Deng, Yunlong Cai, Xiaolong Rao, Yuxin Yin, and Long Rong. 2020. “Plasma Proteomics-Based Identification of Novel Biomarkers in Early Gastric Cancer.” Clinical Biochemistry 76 (February): 5–10. https://doi.org/10.1016/j.clinbiochem.2019.11.001.

Ziani, Paola Rampelotto, Jacson Gabriel Feiten, Jéferson Ferraz Goularte, Rafael Colombo, Bárbara Antqueviezc, Luiza Paul Géa, and Adriane Ribeiro Rosa. 2022. “Potential Candidates for Biomarkers in Bipolar Disorder: A Proteomic Approach through Systems Biology.” Clinical Psychopharmacology and Neuroscience: The Official Scientific Journal of the Korean College of Neuropsychopharmacology 20 (2): 211–27. https://doi.org/10.9758/CPN.2022.20.2.211.

